# Extrinsic MMPs drive epithelial shape change via basal ECM disassembly in the *Drosophila* wing disc

**DOI:** 10.64898/2026.01.21.700823

**Authors:** Chigusa Hinata, Hirotatsu Nakagawa, Shigeaki Nonaka, Katsuya Nozaki, Yoshikatsu Sato, Shizue Ohsawa

## Abstract

Epithelial morphogenesis generates complex tissue architectures with remarkable reproducibility, yet how such self-organized shape transformations are triggered *in vivo* remains poorly understood. Here, we show that epithelial morphogenesis during early pupal development of the *Drosophila* wing disc is regulated non-autonomously through remodeling of the basal extracellular matrix (ECM) by neighboring tissues. Live imaging analyses reveal a previously unrecognized epithelial shape transition at the larval-to-pupal stage, in which the wing disc epithelium transforms from a concave to a convex configuration. Rather than being driven autonomously by the epithelium, this morphogenetic transition depends on adjacent non-epithelial cell populations, including myoblasts and tracheal cells. Mechanistically, systemic ecdysone signaling activates Mmp1 and Mmp2 in these neighboring tissues, leading to spatially restricted basal ECM disassembly that permits collective epithelial shape reorganization. Together, our findings establish ECM remodeling at tissue interfaces as a non-autonomous regulatory layer that enables epithelial self-organization during development.

## Introduction

Tissue morphogenesis generates complex architectures with remarkable reproducibility, a property often attributed to the self-organizing capacity of cells. Accordingly, classical models of morphogenesis have emphasized cell-autonomous behaviors such as cell adhesion, migration, and the generation of contractile forces. In these models, the extracellular matrix (ECM) has traditionally been viewed as a static scaffold that supports cells and passively modulates these processes (Frantz *et al*, 2010). However, accumulating evidence indicates that ECM remodeling, including deposition, reorganization, and localized degradation, plays an active and instructive role in shaping tissue architecture during development (Diaz-de-la-Loza & Stramer, 2024; Matsubayashi, 2022; Sherwood, 2021; Walma & Yamada, 2020). Despite this progress, how ECM remodeling is spatiotemporally coordinated among distinct cell populations to trigger reproducible morphogenetic events *in vivo* remains poorly understood.

In epithelial morphogenesis, interactions between epithelial tissues and surrounding cell types have emerged as key contributors to tissue shaping. For example, during mammalian tooth development, reciprocal paracrine signaling between oral epithelium and neural crest–derived mesenchyme drives progressive epithelial morphogenesis (Hermans *et al*, 2021; Jernvall & Thesleff, 2000). Similar principles operate in other systems, including the *Drosophila* tracheal system, where epithelial branching is guided by positional cues derived from surrounding tissues (Casanova, 2007). Likewise, in mammalian lung development, FGF10 secreted by mesenchymal cells directs epithelial bud outgrowth and branching (Bellusci *et al*, 1997; El Agha & Thannickal, 2023). In addition to paracrine signaling, direct physical interactions with neighboring cells can also instruct epithelial morphogenesis, as demonstrated by recent studies showing that contractile fibroblasts are required for epithelial branching in mammary organoids (Sumbal *et al*, 2024). While these studies highlight the importance of inter-tissue communication, how such interactions are integrated with ECM remodeling to initiate and spatially pattern epithelial self-organization remains largely unresolved.

The *Drosophila* wing disc provides a powerful model to investigate how systemic signals and inter-tissue interactions converge to regulate epithelial morphogenesis. The wing disc is a sac-like epithelial tissue composed of two layers, the disc proper (DP) and the peripodial membrane (PM). At the onset of pupal development, a systemic pulse of the steroid hormone ecdysone triggers extensive tissue remodeling within the wing disc. During this transition, the DP undergoes large-scale, region-specific shape changes that form the adult wing and thoracic structures, whereas the PM detaches and degenerates (Beira & Paro, 2016). Notably, epithelial shape transformation at this developmental stage occurs in close temporal and spatial association with dynamic behaviors of non-epithelial cell populations positioned along the basal side of the DP. Tracheal cells remodel their branching architecture, and myoblasts collectively migrate across the tissue surface during pupal development (Gildor *et al*, 2012; Hayashi & Kondo, 2018). However, how these concurrent epithelial and non-epithelial events are coordinated in space and time to produce reproducible epithelial morphogenesis remains unclear.

Here, we show that epithelial morphogenesis in the *Drosophila* wing disc is instructed through inter-tissue interactions that act to locally remodel the basal ECM. Systemic ecdysone signaling activates MMP activity in neighboring myoblasts and tracheal cells, leading to spatially restricted basal ECM disassembly that permits a concave-to-convex shape transition of the DP epithelium. These findings reveal how systemic developmental cues are translated into localized morphogenetic signals at tissue interfaces, defining a non-autonomous mode of epithelial self-organization during development.

## Results

### The wing disc undergoes a concave-to-convex shape transition during the larval-to-pupal stage

We examined the dynamic process of wing disc morphogenesis during the larval-to-pupal transition using an *ex vivo* culture system (Aldaz *et al*, 2010). Confocal live imaging showed that the wing disc initially bends downward into a concave shape (as indicated by orange arrowheads in the middle panel of Fig. 1I; Video S1). Light-sheet microscopy further revealed that, at the concave stage, the wing disc adopts a ‘Mexican hat’ configuration, where the pouch region forms a prominent central protrusion and the notum and hinge form the surrounding rim structure (schematically illustrated in Fig. 1A; shown in live imaging snapshots in the left panel of Fig. 1B). Remarkably, this ‘Mexican hat’-like concave structure undergoes a transition into a convex shape upon ecdysone stimulation (Fig. B; Video S2). The transition from concave to convex was also observed by confocal microscopy (Fig. 1I, I’; Video S1), although the ‘Mexican hat’-like shape itself at the concave stage was not clearly resolved due to the lower z-resolution of confocal imaging.

**Fig 1.**
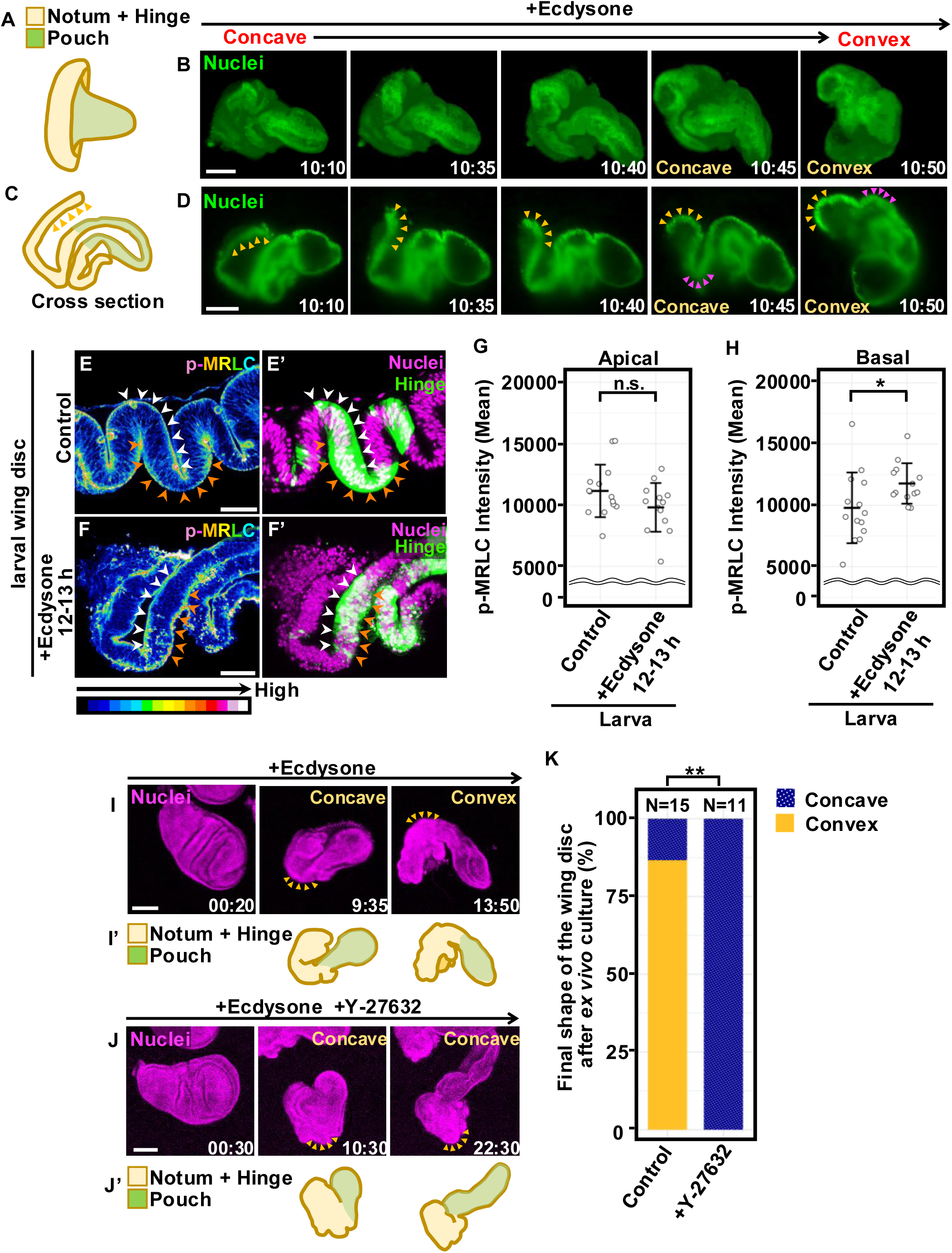
The wing disc undergoes a concave-to-convex shape transition during the larval-to-pupal stage. **(A–D)** Schematic diagrams illustrate the concave three-dimensional shape of the larval wing disc (“Mexican-hat–like” shape) (A) and the corresponding x–z cross-section (C). Time-lapse light-sheet imaging was initiated once the wing discs adopted a concave shape after ecdysone treatment, allowing visualization of the subsequent transition toward a convex shape (B) and the corresponding x–z cross-section (D). Magenta arrowheads indicate the same anatomical position in the disc before and after the curvature change, marking the region that undergoes the concave-to-convex transition. Orange arrowheads indicate a hook-like structure formed prior to this transition. Time is shown as hours:minutes (h:min) from the onset of ecdysone treatment. Scale bar, 100 μm. **(E–F’)** Wing discs in which the hinge region was labeled with *30A-Gal4*-driven GFP (green), either dissected at the third-instar wandering larval stage (E–E’) or dissected and cultured for 12–13 h in the presence of ecdysone (F–F’), were stained with anti-GFP and anti-p-MRLC (pseudocolor) antibodies, together with DAPI (magenta). White and orange arrowheads indicate the apical and basal hinge regions, respectively. Scale bar, 30 μm. **(G–H)** Dot plots showing p-MRLC intensity in the hinge region marked by *30A-Gal4*-driven GFP, with horizontal bars indicating the mean ± standard deviation (SD), corresponding to the representative images shown in (E) (n = 14) and (F) (n = 13). The apical data are shown in (G), and the basal data are shown in (H). *p<0.05, n.s., not significant; Wilcoxon rank-sum test. **(I–J’)** Time-lapse confocal imaging of the ecdysone-treated wing discs carrying *ubi-RFP* (magenta) in the presence of either distilled water (DW) (I) or the ROCK inhibitor Y-27632 (J). Orange arrowheads indicate the same anatomical position in the disc before and after the curvature change, marking the region that normally undergoes the concave-to-convex transition. Schematic diagrams (I’ and J’) illustrate the concave and convex shapes corresponding to the representative images shown in (I) and (J). Time is shown as h:min from the onset of ecdysone treatment. Scale bar, 100 μm. **(K)** Quantification of the final wing disc shape, shown as the percentage of discs that remained concave or transitioned to a convex shape, corresponding to the representative images shown in (I) (n = 15) and (J) (n = 11). **p<0.01; Fisher’s exact test.

Additionally, cross-sectional views of the shape transition, visualized by light-sheet microscopy, showed that the edge region of the wing disc at the concave stage gradually develops a hook-like structure (as indicated by orange arrows in Fig. 1C, D), followed by a rapid transition of the epithelial sheet from concave-downward to convex-upward within five minutes (Fig. 1D). These observations highlight the dynamic nature of epithelial morphogenesis in the wing disc, with a swift transition from a concave to a convex shape in response to ecdysone stimulation. This transient concave-to-convex sequence is considered to reflect a partial eversion process *in vivo* (Pastor-Pareja *et al*, 2004; Tripathi & Irvine, 2022) (Fig. S1A). Importantly, the transition of the wing disc from concave to convex during this process likely determines the final orientation of the adult wing relative to the body surface (Fig. S1A), whereas a disc that remains concave would be expected to adopt the opposite orientation.

To gain further insight into this dynamic process, we focused on the hinge region undergoing the concave-to-convex transition, marked by GFP under the control of the hinge-specific *30A-Gal4* driver (Simmonds *et al*, 1995). Non-muscle myosin II, a major contractile motor during epithelial morphogenesis (Coravos & Martin, 2016; Quintanilla *et al*, 2023), exhibited a marked increase in phosphorylation of its regulatory light chain (p-MRLC) at the basal surface of the hinge region in wing discs at a stage corresponding to the concave-to-convex transition in *ex vivo* culture, compared with the pre-concave larval stage (Fig. 1E–F’; orange arrowheads, quantified in Fig. 1H). In contrast, apical p-MRLC levels remained largely unchanged (Fig. 1E–F’; white arrowheads, quantified in Fig. 1G). Functionally, treatment with the ROCK inhibitor Y-27632 significantly suppressed the concave-to-convex transition (Fig. 1J, J’, quantified in Fig. 1K; Video S3), supporting the idea that this dynamic epithelial morphogenesis is actively driven by ROCK-mediated myosin II contractility.

### MMP-dependent disassembly of the basal ECM is required for the epithelial concave-to-convex shape transition

We next sought to identify the key factor required for the epithelial concave-to-convex transition in the wing disc. ECM remodeling has been reported to occur during early pupal morphogenesis, including in the wing disc (De Las Heras *et al*, 2018; Diaz-de-la-Loza *et al*, 2020; Diaz-de-la-Loza *et al*, 2018; Srivastava *et al*, 2007; Sun *et al*, 2021; Tsuboi *et al*, 2023). Indeed, we observed a progressive decrease in Collagen IV, the main component of the basal ECM (α2-subunit encoded by *viking*), not only in the PM but also in the DP epithelium of pupal wing discs (Fig. S2A–D’’), consistent with previous findings (Diaz-de-la-Loza *et al*., 2020; Diaz-de-la-Loza *et al*., 2018). A similar decrease was also detected in the DP of *ex vivo* cultured discs stimulated with ecdysone (Fig. 2A–B’’, quantified in Fig. 2E). This decrease in Collagen IV was largely mediated by MMP activity, as treatment with the broad-spectrum MMP inhibitor GM6001 significantly suppressed the reduction in Collagen IV levels in the DP under ecdysone stimulation (Fig. 2C–D’’, quantified in Fig. 2E).

**Fig 2.**
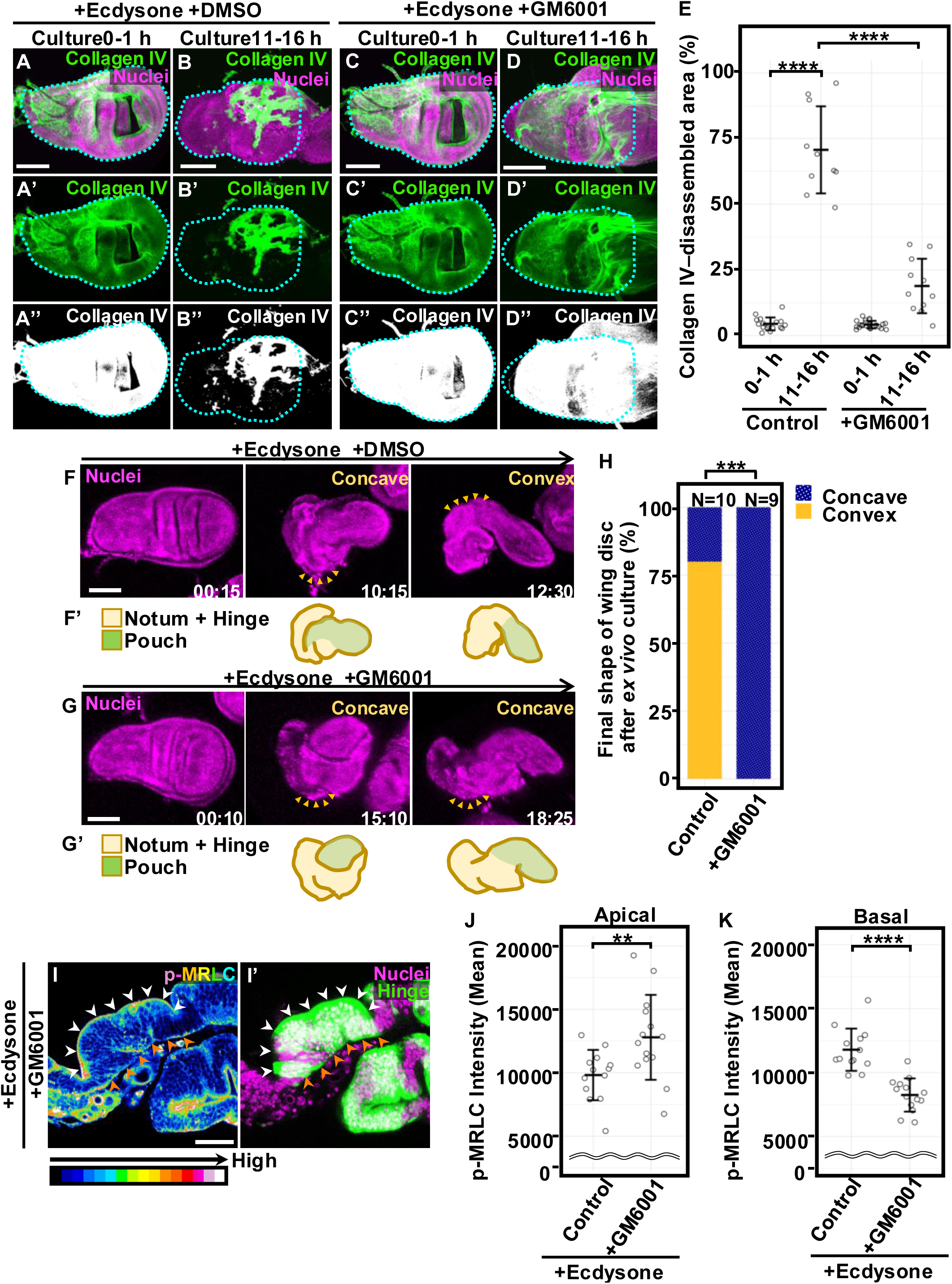
MMP-dependent disassembly of the basal ECM is required for the epithelial concave-to-convex shape transition. **(A–D’’)** Wing discs carrying *viking::GFP* (green) and *ubi-RFP* (magenta) were dissected and treated with ecdysone for 0–1 h or 11–16 h in the presence of either DMSO (A–B’’) or the MMP inhibitor GM6001 (C–D’’). The black-and-white images (A’’), (B’’), (C’’), and (D’’) represent binarized versions of (A’), (B’), (C’), and (D’), respectively. Scale bar, 100 μm. **(E)** Dot plots showing the percentage of the Collagen IV-disassembled area on the basal side, with horizontal bars indicating the mean ± SD, corresponding to the representative images shown in (A’’) (n = 17), (B’’) (n = 10), (C’’) (n = 22), and (D’’) (n = 11). ****p<0.0001; pairwise comparisons using the Wilcoxon rank-sum test with Bonferroni correction. **(F–G’)** Time-lapse confocal imaging of the ecdysone-treated wing discs carrying *ubi-RFP* (magenta) in the presence of either DMSO (F) or the MMP inhibitor GM6001 (G). Orange arrowheads indicate the same anatomical position in the disc before and after the curvature change, marking the region that normally undergoes the concave-to-convex transition. Schematic diagrams (F’ and G’) illustrate the concave and convex shapes corresponding to the representative images shown in (F) and (G). Time is shown as h:min from the onset of ecdysone treatment. Scale bar, 100 μm. **(H)** Quantification of the final wing disc shape, shown as the percentage of discs that remained concave or transitioned to a convex shape, corresponding to the representative images shown in (F) (n = 10), (G) (n = 9). ***p<0.001; Fisher’s exact test. **(I–I’)** Wing discs in which the hinge region was marked by *30A-Gal4*-driven GFP (green) were dissected, treated with ecdysone in the presence of the MMP inhibitor GM6001 (F–F’), and stained with anti-GFP and anti–p-MRLC (pseudocolor) antibodies, together with DAPI (magenta). White and orange arrowheads indicate the apical and basal hinge regions, respectively. Scale bar, 30 μm. **(J–K)** Dot plots showing p-MRLC intensity in the hinge region marked by *30A-Gal4*-driven GFP, with horizontal bars indicating the mean ± SD, corresponding to the representative images shown in Figure 1F (n = 13) and Figure 2I (n = 14). Apical and basal data are shown in (J) and (K), respectively. **p<0.01, ****p<0.0001; Wilcoxon rank-sum test.

We next examined whether MMP activity contributes to the concave-to-convex epithelial shape transition of the wing disc. In *ex vivo* cultures, treatment with the MMP inhibitor markedly suppressed the ecdysone-induced transition (Fig. 2G, G’, compared with Fig. 2F, F’; quantified in Fig. 2H; Video S5, compared with Video S4), suggesting that MMP activity is important for inducing this morphological transition. Consistently, pharmacological inhibition of MMP activity markedly reduced the basal increase in p-MRLC levels in the hinge region, which was marked by *30A-Gal4*-driven GFP (Fig. 2I, I’; orange arrowheads, compared with Fig.1F, F’, quantified in Fig. 2K). At the same time, apical p-MRLC levels were aberrantly elevated (Fig. 2I, I’; white arrowheads, compared with Fig.1F, F’, quantified in Fig. 2J). This supports the idea that MMP activity could coordinate the spatial activation of myosin II during this process. Importantly, treatment with the ROCK inhibitor Y-27632 did not prevent the ecdysone-induced decrease in Collagen IV levels in wing discs (Fig. S1D–E’’, compared with Fig. S1B-C’’, quantified in Fig. S1F). Together, these results suggest that MMP-dependent Collagen IV disassembly acts upstream of ROCK-mediated myosin contractility and the epithelial concave-to-convex shape transition.

### Myoblasts and tracheal cells are spatially associated with localized basal ECM disassembly during the larval-to-pupal transition

Given that MMP activity is required for the epithelial shape transition, we next examined how ECM disassembly downstream of MMP activity is spatially and temporally organized within the wing disc by performing time-lapse imaging on *ex vivo* cultures embedded in agarose gel to mechanically restrict tissue movement. We found that the reduction of Collagen IV in the DP was not uniform in the wing disc under ecdysone stimulation. Instead, localized holes appeared in the Collagen IV sheets, initiating a progressive peeling process as these openings expanded (Fig. 3A, A’; Video S6). A similar pattern was observed for Trol (the *Drosophila* homolog of Perlecan), which exhibited localized perforation followed by expansion and peeling (Fig. S3A, A’; Video S7), suggesting that such localized basal ECM disassembly may be a general feature of matrix remodeling in the wing disc during the larval-to-pupal transition.

**Fig 3.**
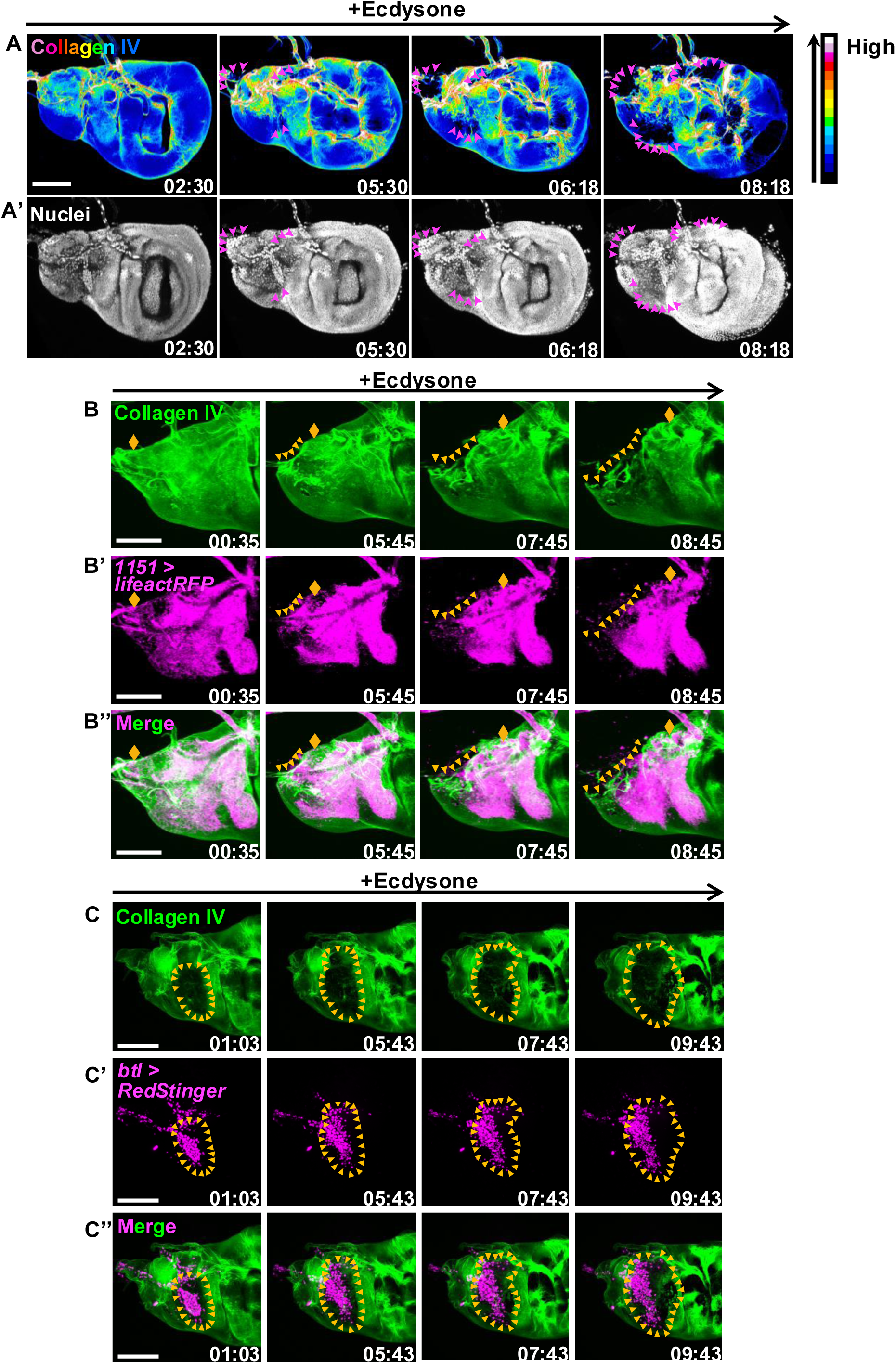
Myoblasts and tracheal cells are spatially associated with localized basal ECM disassembly during the larval-to-pupal transition. **(A–A’)** Time-lapse confocal imaging of an ecdysone-treated wing disc carrying *viking::GFP* (pseudocolor) and *ubi-RFP* (white). Magenta arrowheads indicate regions where Collagen IV has undergone disassembly in the notum and hinge regions. Time is shown as h:min from the onset of ecdysone treatment. Scale bar, 100 µm. **(B–B’’)** Time-lapse confocal imaging of an ecdysone-treated wing disc carrying *viking::GFP* (green) and expressing *lifeactRFP* (magenta) under the control of *1151-Gal4*. Orange arrowheads indicate regions of Collagen IV disassembly associated with myoblast migration. The orange rhombus denotes a reference point within the migrating myoblast cluster. Time is shown as h:min from the onset of ecdysone treatment. Scale bar, 100 µm. **(C–C’’)** Time-lapse confocal imaging of an ecdysone-treated wing disc carrying *viking::GFP* (green) and expressing *RedStinger* (magenta) under the control of *btl-Gal4*. Orange arrowheads indicate the regions of Collagen IV-disassembled area around the trachea cells. Time is shown as h:min from the onset of ecdysone treatment. Scale bar, 100 µm.

To further investigate the gradual decrease of Collagen IV in the DP during the ecdysone-induced morphological transition of the wing disc, we developed a new live imaging method that enables visualization of the x–z optical sections in *ex vivo* cultured wing discs using confocal microscopy. Unexpectedly, we observed cells in the notum region located outside the DP migrating and apparently disrupting Collagen IV network (Fig. S3B, B’; Video S8). These observations imply that Collagen IV disassembly in this region is driven by active cellular regulation rather than by passive matrix dynamics.

Myoblasts and tracheal cells in the notum region of wing discs, positioned between the basal surface of the DP and the basal ECM (Fig. S3C–H’’), undergo dynamic morphological changes during pupal development. Myoblasts migrate toward the nascent dorsal longitudinal muscle (DLM) myotubes, which are derived from a persistent set of larval muscles (Fernandes *et al*, 1991), while tracheal cells exhibit growth and branching morphogenesis (Guha & Kornberg, 2005). Time-lapse imaging of *ex vivo* cultured wing discs revealed that both myoblasts and tracheal cells, labeled with the myoblast-specific *1151-Gal4* driver (Roy & VijayRaghavan, 1997) and the pan-tracheal *breathless (btl)-Gal4* driver (Shiga *et al*, 1996) respectively, are actively mobile under ecdysone stimulation (Video S9, S10). Notably, the onset of collective myoblast migration and tracheal cell activity coincided spatially and temporally with local Collagen IV disassembly (Fig. 3B–3C’’). Together, these observations suggest that myoblasts and tracheal cells may contribute to basal ECM disassembly during the ecdysone-induced concave-to-convex transition of wing discs.

### Mmp1 and Mmp2 in myoblasts and tracheal cells drive Collagen IV disassembly and epithelial shape transition

We next investigated whether the *Drosophila* MMPs, Mmp1 and Mmp2, are responsible for the observed ECM disassembly associated with myoblasts and tracheal cells. To monitor their expression, we used an *mmp1-GFP* transcriptional reporter (Wang *et al*, 2010) and an *Mmp2::GFP* knock-in reporter tagging the C-terminus of the Mmp2 RB isoform (Diaz-de-la-Loza *et al*., 2020). Using these reporters, we found that both Mmp1 and Mmp2 were upregulated in a subset of myoblasts marked by *1151-Gal4*-driven RedStinger in pupal wing discs at 1–1.5 h after puparium formation (APF), compared with third-instar wandering larvae (Mmp1: Fig. 4A–A’’, compared with Fig. S4A–A’’; Mmp2: Fig. 4C–C’’, compared with Fig. S4C–C’’). Mmp2 was likewise upregulated in tracheal cells in pupal wing discs at 1–1.5 h APF, compared with third-instar wandering larvae, with the tracheal lumen visualized by anti-Gasp antibody staining (Tsarouhas *et al*, 2007) (Fig. 4D–D″ and Fig. S4D–D″). In addition, Mmp1 was already strongly expressed in tracheal cells, marked by *btl-Gal4*-driven RedStinger, from the third-instar wandering larval stage through 1–1.5 h APF (Fig. 4B–B’’ and Fig. S4B–B’’).

**Fig 4.**
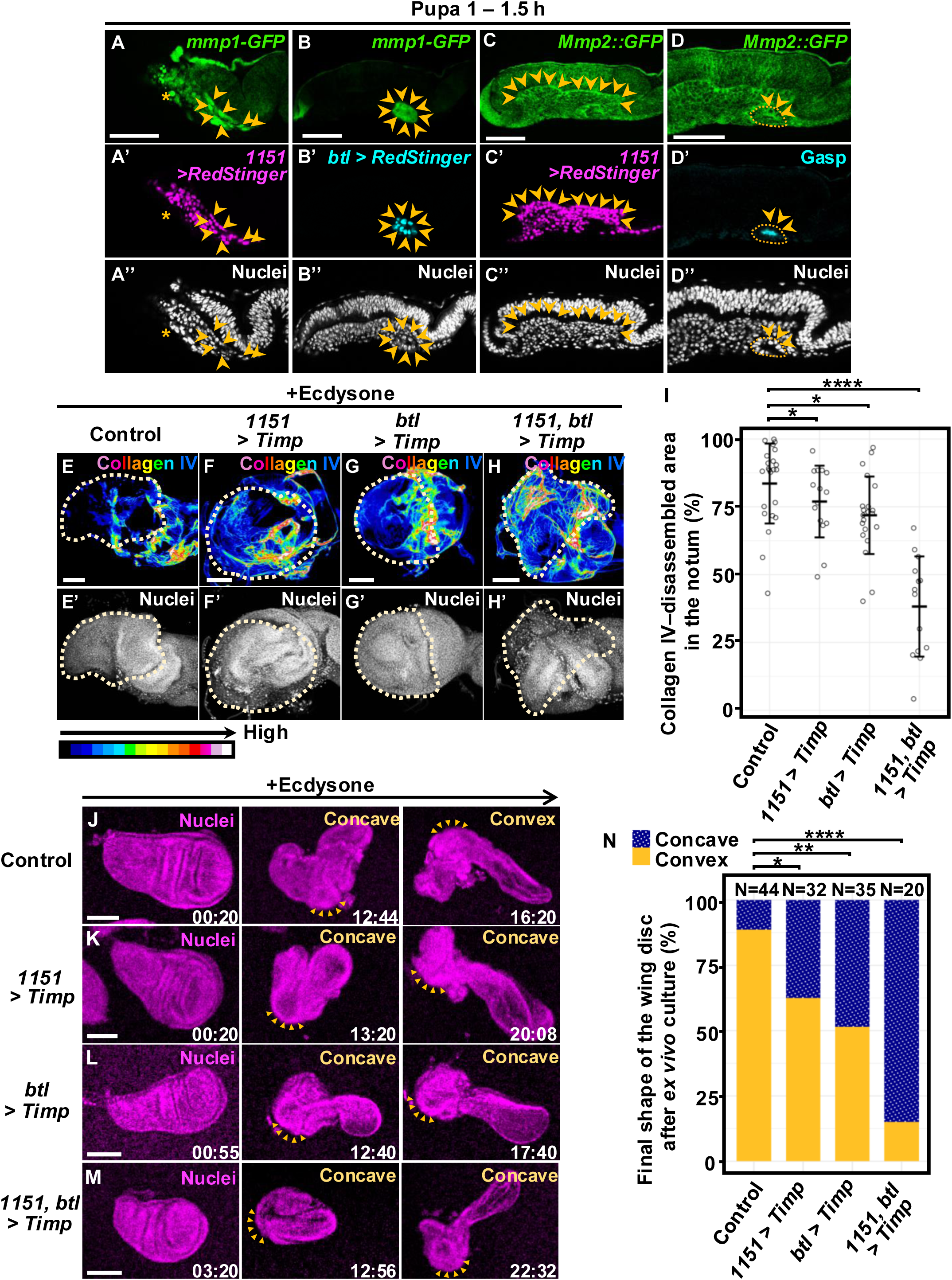
Mmp1 and Mmp2 in myoblasts and tracheal cells drive Collagen IV disassembly and the epithelial shape transition. **(A–B’’)** Wing discs carrying *mmp1-GFP* transcriptional reporter (green) and expressing *RedStinger* under the control of *1151-Gal4* (magenta) or *btl-Gal4* (cyan) were dissected at 1–1.5 h after puparium formation (APF) and stained with anti-GFP antibody and DAPI (white). Orange arrowheads indicate Mmp1-positive myoblasts or tracheal cells. An asterisk indicates a Mmp1-positive tracheal cell shown in (A). Scale bar, 50 μm. **(C–C’’)** A wing disc carrying the *Mmp2::GFP* knock-in reporter (green) and expressing *RedStinger* under the control of *1151-Gal4* (magenta) at 1–1.5 h APF, stained with anti-GFP antibody and DAPI (white). Orange arrowheads indicate Mmp2-positive myoblasts. Scale bar, 50 μm. **(D–D’’)** A wing disc carrying the *Mmp2::GFP* knock-in reporter (green) at 1–1.5 h APF, stained with anti-GFP and anti-Gasp (cyan) antibodies, together with DAPI (white). Orange dotted lines outline the tracheal tube. Orange arrowheads indicate Mmp1-positive tracheal cells. Scale bar, 50 μm. **(E–H’)** Wing discs carrying *viking::GFP* (pseudocolor) were dissected, treated with ecdysone, and stained with anti-GFP antibody and DAPI (white). Images are shown for control (E) and for discs expressing Timp under the control of *1151-Gal4* (F), *btl-Gal4* (G), or both *1151-Gal4* and *btl-Gal4* (H). Yellow dotted lines outline the notum regions. Scale bar, 50 μm. **(I)** Dot plots showing the percentage of Collagen IV-disassembled area in the notum, where Collagen IV signals could be more reliably quantified than in the hinge region, with horizontal bars indicating the mean ± SD, corresponding to the representative images shown in (E) (n = 23), (F) (n = 15), (G) (n = 21), and (H) (n = 14). *p<0.05, ****p<0.0001; pairwise comparisons using the Wilcoxon rank-sum test with BH correction. **(J–M)** Time-lapse confocal imaging of the ecdysone-treated wing discs carrying *ubi-RFP* (magenta). Images are shown for control (J) and for discs expressing Timp under the control of *1151-Gal4* (K), *btl-Gal4* (L), or both *1151-Gal4* and *btl-Gal4* (M). Orange arrowheads indicate the same anatomical position before and after the curvature change, marking the region that normally undergoes the concave-to-convex transition. Time is shown as h:min from the onset of ecdysone treatment. Scale bar, 100 μm. **(N)** Quantification of the final wing disc shape, shown as the percentage of discs that remained concave or transitioned to a convex shape, corresponding to the representative images shown in (J) (n = 44), (K) (n = 32), (L) (n = 35), and (M) (n = 20). *p<0.05, **p<0.01, ****p<0.0001; Fisher’s exact test with Holm correction.

To functionally test the role of MMPs in these cell types, we overexpressed Timp (tissue inhibitor of metalloproteinases) (Godenschwege *et al*, 2000), in myoblasts and tracheal cells using the *1151-Gal4* and *btl-Gal4* drivers, respectively. Dual overexpression of Timp in both myoblasts and tracheal cells markedly suppressed the reduction of Collagen IV in the dorsal epithelium, including the notum and hinge regions, of *ex vivo* cultured wing discs under ecdysone stimulation (Fig. 4H, H’, compared with Fig. 4E, E’, quantified in Fig. 4I), whereas Collagen IV disassembly in the pouch region remained unaffected (Fig. S4H, H’, compared with Fig. S4E, E’, quantified in Fig. S4I). Light-sheet microscopy further revealed that, at the concave stage, Collagen IV remained tightly associated with the dorsal epithelium upon overexpression of Timp in both myoblasts and tracheal cells (Fig. S4K–K’’, compared with Fig. S4J-J’’; Video S11, S12). Strikingly, both light-sheet and confocal time-lapse imaging showed that this manipulation strongly inhibited the concave-to-convex transition of the wing disc under the same conditions (Fig. 4M, compared with Fig. 4J, quantified in Fig. 4N; Video S12, S13). Subsequent confocal analyses revealed that overexpression of Timp in either myoblasts or tracheal cells alone also partially suppressed both Collagen IV disassembly and the concave-to-convex transition of the wing disc, although the effects were weaker than those observed upon dual overexpression (Fig. 4F–G’, quantified in Fig. 4I; Fig. 4K, L, quantified in Fig. 4N; Video S14, S15). These results demonstrate that MMP activity in myoblasts and tracheal cells is functionally required for Collagen IV disassembly and the ensuing epithelial morphogenesis.

We next examined which MMPs were responsible for these effects. Knockdown of *Mmp1* or *Mmp2* individually in both cell types partially suppressed the reduction of Collagen IV in the dorsal epithelium (Fig. S4M–N’, compared with Fig. S4L, L’, quantified in Fig. S4P), whereas simultaneous knockdown of both genes resulted in a much stronger suppression (Fig. S4O, O’, quantified in Fig. S4P). Similarly, the concave-to-convex transition of the wing disc was inhibited by either single knockdown, with a more pronounced effect upon simultaneous depletion of both MMPs (Fig. S4R–T’, compared with Fig. S4Q, Q’, quantified in Fig. S4U). These data suggest that Mmp1 and Mmp2 in myoblasts and tracheal cells act cooperatively to mediate Collagen IV disassembly, thereby driving the epithelial shape transition of the wing disc.

### Ecdysone signaling functions upstream of Mmp1 and Mmp2 to drive the epithelial shape transition

It has been reported that *Mmp1* and *Mmp2* are transcriptionally upregulated downstream of ecdysone signaling during the larval-to-pupal transition (Diaz-de-la-Loza *et al*., 2018; Fraire-Zamora *et al*, 2021; Graveley *et al*, 2011; mod *et al*, 2010; Song *et al*, 2025). To determine whether ecdysone signaling regulates MMP-mediated Collagen IV disassembly in myoblasts and tracheal cells, we simultaneously knocked down the *ecdysone receptor* (*EcR*) in these cells using the *1151-Gal4* and *btl-Gal4* drivers. Larvae were cultured for 63–75 h at 18 °C to avoid embryonic lethality before shifting to 30 °C for *RNAi* induction (Fig. S5A). Under the dual *EcR* knockdown conditions, tracheal cell viability was severely compromised, as assessed by anti-Gasp antibody staining, thereby precluding an accurate assessment of Mmp1 and Mmp2 expression (Fig. S5B–D’). In contrast, under the same dual-knockdown conditions, myoblasts, identified by anti-Zfh1 antibody staining (Lai *et al*, 1991), showed markedly reduced expression of Mmp1 (Fig. 5C–D’, compared with Fig. 5A–B’, quantified in Fig. 5E) and Mmp2 (Fig. 5G, G’, compared with Fig. 5F, F’, quantified in Fig. 5H), detected using an anti-Mmp1 antibody and the *Mmp2::GFP* reporter, respectively. These reductions in Mmp1 and Mmp2 expression were accompanied by a strong suppression of Collagen IV disassembly in the dorsal epithelium (Fig. 5J, J’, compared with Fig. 5I, I’, quantified in Fig. 5K) and a concomitant inhibition of the concave-to-convex transition of the wing disc (Fig. 5M–N’, compared with Fig. 5L, L’, quantified in Fig. 5O).

**Fig 5.**
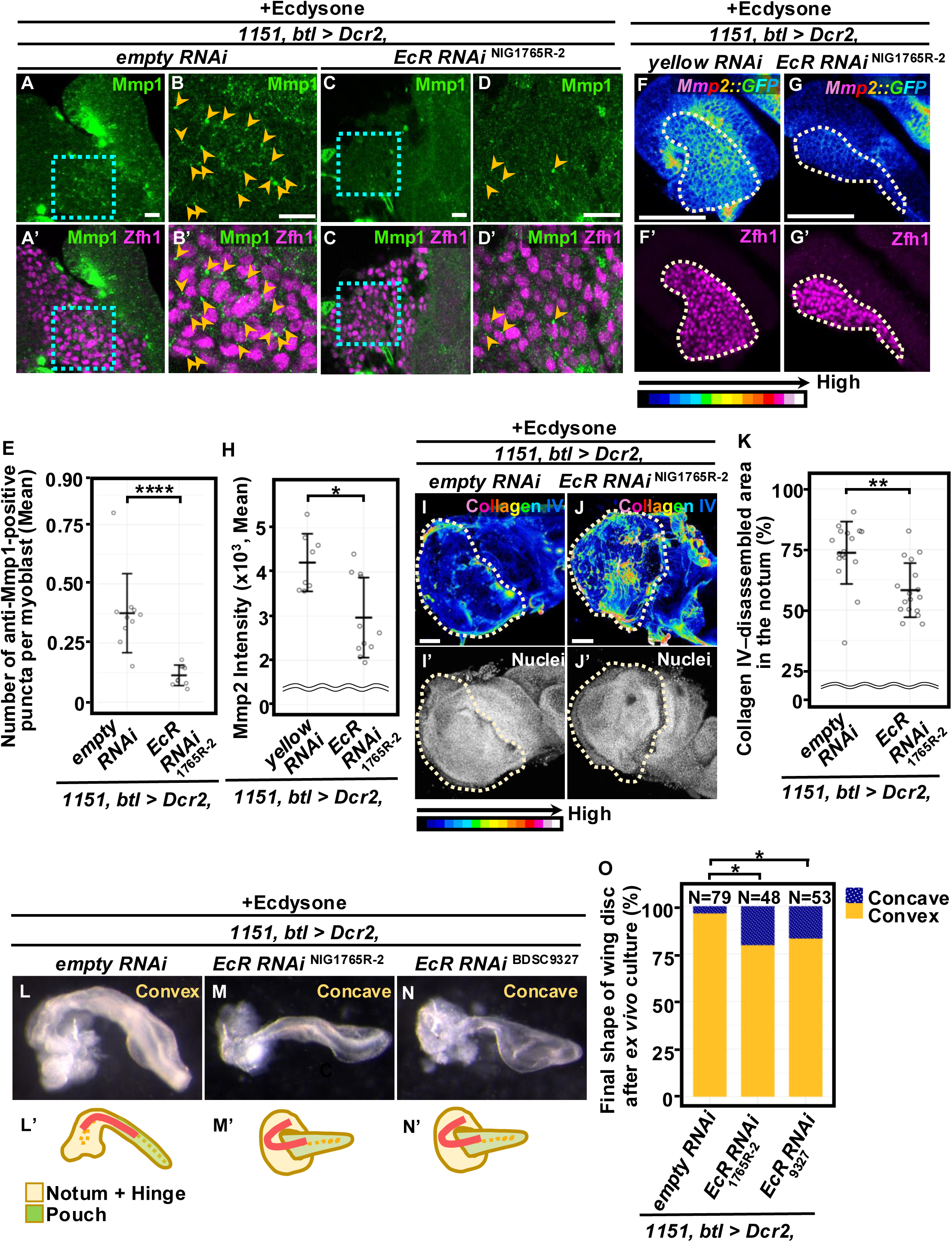
Ecdysone signaling functions upstream of Mmp1 and Mmp2 to drive the epithelial shape transition. **(A–D’’)** Wing discs were dissected, treated with ecdysone, and stained with anti-Mmp1 (green) and anti-Zfh1 (magenta) antibodies. Images are shown for discs expressing *empty-RNAi* (VDRC60100) (A–B’) or *EcR-RNAi* (NIG1765R-2) (C–D’), together with Dicer2 (Dcr2), under the control of *1151-Gal4* and *btl-Gal4*. (B–B’) and (D–D’) show magnified views of the regions outlined by cyan dotted lines in (A–A’) and (C–C’), respectively. Larvae carrying *Tub-Gal80^ts^* were maintained under the temperature conditions illustrated in Fig. S5A. Orange arrowheads indicate the anti-Mmp1-positive puncta surrounding Zfh1-positive myoblast nuclei. Scale bar, 10 μm. **(E)** Dot plots showing the mean number of anti-Mmp1-positive puncta per myoblast, with horizontal bars indicating the mean ± SD, corresponding to the representative images shown in (A–B) (n = 10) and (C–D) (n = 9). ****p<0.0001; Wilcoxon rank-sum test. **(F–G’)** Wing discs carrying *Mmp2::GFP* knock-in reporter (pseudocolor) were dissected, treated with ecdysone, and stained with anti-GFP and anti-Zfh1 (magenta) antibodies. Images are shown for discs expressing *yellow-RNAi* (NIG3757R-1) (F–F’) or *EcR-RNAi* (NIG1765R-2) (G–G’), together with Dcr2, under the control of *1151-Gal4* and *btl-Gal4*. Larvae bearing *Tub-Gal80^ts^* were maintained under the temperature conditions indicated in Fig. S5A. Scale bar, 50 μm. **(H)** Dot plots showing the mean intensity of *Mmp2::GFP* knock-in reporter in myoblasts, with horizontal bars indicating the mean ± SD, corresponding to the representative images shown in (F) (n = 8) and (G) (n = 11). *p<0.05; Wilcoxon rank-sum test. **(I–J’)** Wing discs were dissected, treated with ecdysone, and stained with anti-Collagen IV antibody (pseudocolor) and DAPI (white). Images are shown for discs expressing *empty-RNAi* (VDRC60100) (I) or *EcR-RNAi* (NIG1765R-2) (J), together with Dcr2, under the control of *1151-Gal4* and *btl-Gal4*. Larvae carrying *Tub-Gal80^ts^* were maintained under the temperature conditions illustrated in Fig. S5A. Scale bar, 50 μm. **(K)** Dot plots showing the percentage of the Collagen IV-disassembled area in the notum, with horizontal bars indicating the mean ± SD, corresponding to the representative images shown in (I) (n = 17) and (J) (n = 17). **p<0.01; Wilcoxon rank-sum test. **(L–N’)** Wing discs were dissected and treated with ecdysone for 24 h. Images are shown for discs expressing *empty-RNAi* (VDRC60100) (L), *EcR-RNAi* (NIG1765R-2) (M), or *EcR-RNAi* (BDSC9327) (N), together with Dcr2, under the control of *1151-Gal4* and *btl-Gal4*. Larvae carrying *Tub-Gal80^ts^* were maintained under the temperature conditions illustrated in Fig. S5A. Schematic diagrams (L’, M’ and N’) illustrate the concave and convex shapes corresponding to the representative images shown in (L), (M), and (N). **(O)** Quantification of the final wing disc shape, shown as the percentage of discs that remained concave or transitioned to a convex shape, corresponding to the representative images shown in (L) (n = 79), (M) (n = 48), and (N) (n =53). *p<0.05; Fisher’s exact test with Holm correction.

Together, these findings reveal a regulatory cascade in which ecdysone signaling activates Mmp1 and Mmp2 expression in myoblasts (and likely in tracheal cells as well), thereby promoting localized Collagen IV disassembly that drives the epithelial concave-to-convex transition of the wing disc during the larval-to-pupal morphogenesis.

## Discussion

Here, we identify a non-autonomous mechanism of epithelial morphogenesis in the *Drosophila* wing disc, in which epithelial architecture is instructed by neighboring, non-epithelial cells rather than arising solely from autonomous programs within the epithelium. We further show that this inter-tissue instruction is mediated by ecdysone-driven activation of MMPs in myoblasts and tracheal cells, leading to localized basal ECM disassembly that couples systemic developmental cues to spatially restricted morphogenetic remodeling at tissue interfaces (Fig. 6).

**Fig 6.**
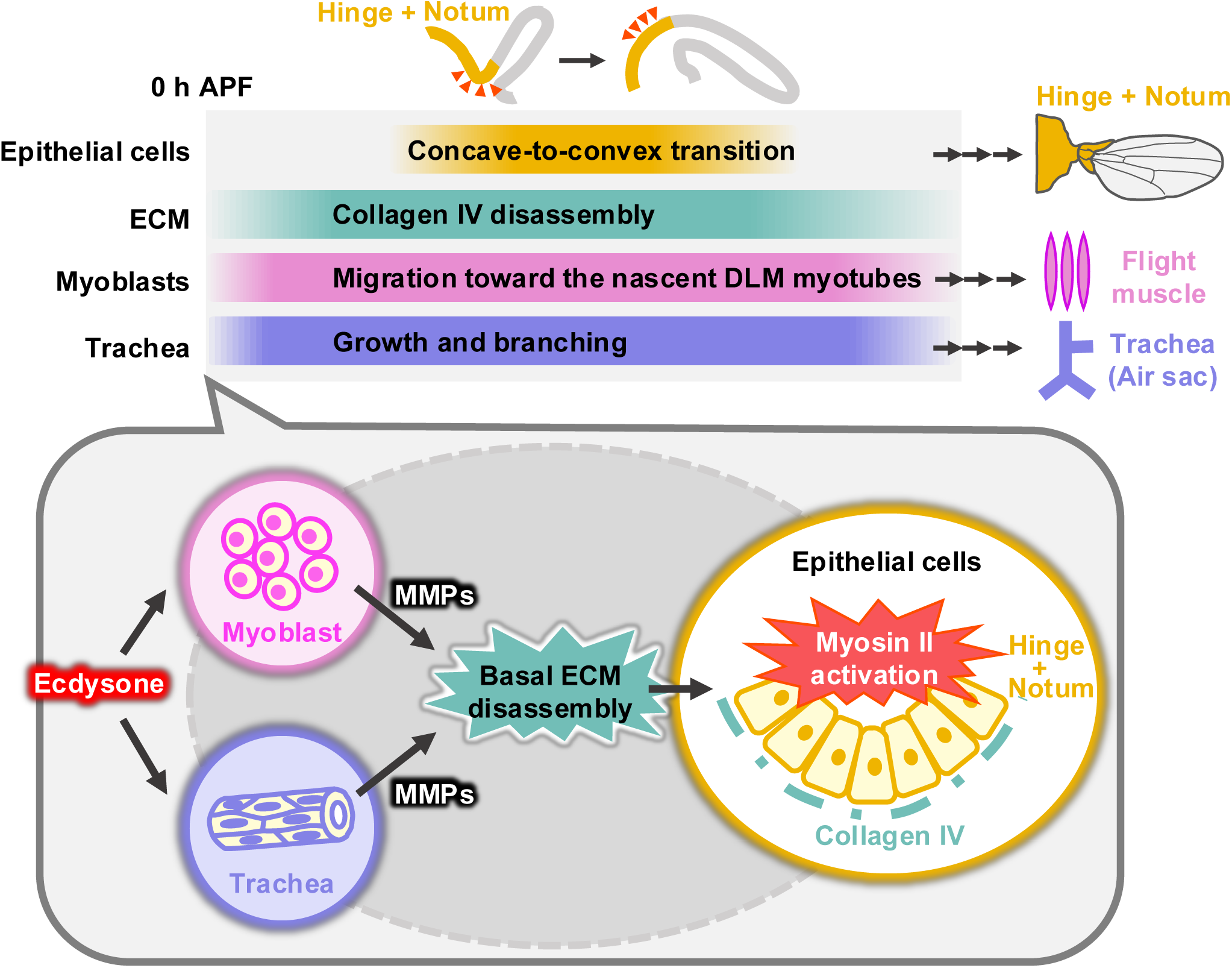
Non-autonomous epithelial concave-to-convex shape transition of the wing disc. Model for the non-autonomous epithelial concave-to-convex shape transition of the wing disc, driven by MMP-dependent ECM disassembly originating from myoblasts and tracheal cells. See text for details.

Our results suggest that basal ECM remodeling during wing disc morphogenesis is regulated in a region-specific manner, with distinct cellular sources of MMP activity operating in different regions: genetic inhibition of MMP activity in myoblasts and tracheal cells selectively suppressed Collagen IV disassembly in the dorsal epithelium, while Collagen IV levels in the wing pouch remained largely unaffected. Consistent with this pattern, it has been shown that MMP activity within the wing pouch is closely associated with epithelial remodeling linked to tissue expansion in this region during the larval-to-pupal transition (Diaz-de-la-Loza et al., 2018). Distinct modes of ECM-mechanical coupling have also been described within the larval wing disc: during fold formation, the central hinge fold is driven by local ECM reduction and the subsequent decrease in basal tension, whereas the fold at the hinge–pouch boundary is generated by cytoskeletal reorganization associated with changes in lateral tension rather than basal ECM reduction (Sui *et al*, 2018). More broadly, organ identity can impose additional constraints on ECM dynamics, as exemplified by the Hox gene *Ultrabithorax*, which suppresses Mmp1 expression in the haltere pouch and delays Collagen IV clearance relative to the wing, thereby restricting appendage shape changes (De Las Heras *et al*., 2018). Together, these observations suggest that ECM remodeling is regulated not only in a region-specific manner within the wing disc, but also in an organ-specific manner across homologous appendages, leading to distinct morphogenetic outcomes.

Our analysis of ROCK-dependent myosin II activation provides a potential mechanistic link between ECM remodeling and epithelial shape change. Pharmacological inhibition of ROCK-dependent myosin contractility blocked the concave-to-convex transition, whereas ECM disassembly, as indicated by Collagen IV reduction, proceeded normally under the same conditions. These observations suggest that MMP-dependent basal ECM remodeling occurs upstream of epithelial contractility and may create a permissive condition for epithelial shape change. This interpretation contrasts with the prevailing paradigm in which increased ECM stiffness or externally applied mechanical forces promote actomyosin contractility through integrin-mediated mechanotransduction (Clarke & Martin, 2021; Humphrey *et al*, 2014). At a broader conceptual level, however, our findings are consistent with previous work showing that local weakening of integrin-dependent basal cell–ECM adhesion can drive epithelial folding through reorganization of the basal actomyosin cytoskeleton in the *Drosophila* wing disc (Valencia-Exposito *et al*, 2025). Nevertheless, whether hinge-localized myosin activation, together with ECM disassembly, is by itself sufficient to drive curvature inversion remains to be determined, raising the possibility that additional tissue-scale mechanical constraints, such as global tissue geometry or forces from surrounding regions, may be required.

Finally, our findings highlight a broader mode of tissue morphogenesis in which interactions between distinct cell populations are coordinated through dynamic ECM remodeling to drive tissue-scale shape changes. For example, ECM disassembly by neighboring tissues has been shown to trigger epithelial proliferation and growth during abdominal epidermis formation in *Drosophila* (Davis *et al*, 2022). Similarly, spatiotemporally regulated basal ECM remodeling generates perforations through MMP activity, facilitating embryo growth and primitive streak extension prior to gastrulation in early mouse embryogenesis (Kyprianou *et al*, 2020). Analogous mechanisms may also operate in pathological settings, where stromal cells remodel the ECM to enable epithelial invasion and expansion in mammalian tumors (Lu *et al*, 2012; Winkler *et al*, 2020). Together, the available evidence suggests that ECM-mediated coordination between distinct cell populations may represent a conserved strategy for regulating tissue morphogenesis across developmental and disease contexts.

## Method

### Fly strains

Fly stocks were cultured at 25°C on standard fly food. When *Tub-Gal80^ts^*was used, flies were maintained at 18 °C until the second-instar larval stage and then shifted to 29°C for 3–5 days. The strains used in this study are as follows: *Tub-Gal4* (BDSC5138), *30A-Gal4* (DGRC106507), *1151-Gal4* (L. S. Shashidhara) (Roy & VijayRaghavan, 1997), *btl-Gal4* (DGRC109128), *vkg::GFP* (DGRC110692), *ubi-RFP* (BDSC35496, BDSC30555), *trol-GFP* (DGRC110836), *mmp1-GFP* (D. Bohmann) (Wang *et al*., 2010), *Mmp2::GFP* (B.J. Thompson) (Diaz-de-la-Loza *et al*., 2020), *UAS-myc-nls-SCAT3* (M. Miura) (Koto *et al*, 2009), *UAS-GFP* (BDSC1521), *UAS-lifeactRFP* (BDSC58362), *UAS-RedStinger* (BDSC8545, BDSC8546, BDSC8547), *UAS-Timp* (J.C. Pastor-Pareja) (Srivastava *et al*., 2007), *UAS-Mmp1-RNAi* (NIG4859R-1), *UAS-Mmp2-RNAi* (NIG1794-1R-2), *UAS-empty-RNAi* (VDRC60100), *UAS-yellow-RNAi* (NIG3757R-1), *UAS-EcR-RNAi* (NIG1765R-2, BDSC9327), *UAS-Dcr2* (BDSC24650), and *Tub-Gal80^ts^*(BDSC7108, BDSC7017).

### Sample preparation for *ex vivo* culture of wing discs

Third-instar wandering larvae were dissected in Schneider’s *Drosophila* Medium (GIBCO, #21720024) supplemented with 2% fetal bovine serum (FBS) and 0.5% penicillin-streptomycin (GIBCO, #15140122). Dissected wing discs were cultured in Schneider’s *Drosophila* Medium supplemented with 2% FBS, 0.5% penicillin-streptomycin, 0.5% methylcellulose (Sigma-Aldrich, #M0387) and 1 µg/ml 20-hydroxyecdysone (Sigma-Aldrich, # H5142), either on glass bottom dishes (Matsunami glass, #D11130H) or on glass slides with a spacer (Grace Bio-labs, #654006), and maintained at 30°C. For ROCK inhibitor treatment experiments (Fig. 1I–K, Fig. S1B–F, Video S2, Video S3), wing discs were mounted in medium supplemented with 400 µM Y-27632 and 0.8% distilled water (FUJIFILM Wako, #253-00513), or in medium containing 0.8% distilled water alone as a control. For MMP inhibitor treatment experiments (Fig. 2A–H, Video S4, Video S5), wing discs were mounted in medium supplemented with 50 µM GM6001 and 0.5% DMSO (Abcam, #ab120845), or in medium containing 0.5% DMSO alone as a control. For observation of ECM disassembly (Fig. 3A–A’, Fig. S3A–A’, Video S6, Video S7) together with the dynamics of myoblasts or tracheal cells (Fig. 3B–C’’, Video S9, Video S10), wing discs were embedded in 1% low-melting agarose gel (Sigma-Aldrich, #A4018) and cultured in medium. For observation of ECM disassembly and cellular dynamics within the wing disc (Fig. S3B–B’, Video S8), a sheet of 3% low-melting agarose gel with a hole approximately the same size as the wing disc was prepared, into which the wing disc was placed and cultured in medium.

### Time-lapse light-sheet imaging of *ex vivo* cultured wing discs

Wing discs used for time-lapse imaging were prepared as described above in the “Sample preparation for *ex vivo* culture of wing discs” section. For light-sheet imaging, an FEP tube was attached to a capillary using UV-curable adhesive. Wing discs were cultured while being supported from below by a layer of 1% low-melting agarose gel, and the assembly was further secured by covering the gel and the base of the tube with wax. Time-lapse imaging was performed at 5-min intervals using a Zeiss Lightsheet Z.1 microscope equipped with a Zeiss W Plan-Apochromat 20x/1.0 NA water-immersion objective. The images of the wing discs were analyzed using Imaris software.

### Time-lapse confocal imaging of *ex vivo* cultured wing discs

Wing discs used for time-lapse imaging were prepared as described above in the “Sample preparation for *ex vivo* culture” section. A coverslip or a dissolved oxygen membrane (YSI, #5793) was placed on top of the spacer, and the sample was then sealed with halocarbon oil (Sigma-Aldrich, #H8898) to prevent dehydration. Time-lapse imaging was performed at 10–15-min intervals using Leica SP5, Leica SPE, and Leica STELLARIS confocal microscopes. On all Leica microscopes, a Leica HC PL APO 10×/0.40 NA objective was used, while the Leica HC PL APO 40×/1.25 NA GLYC objective and the Olympus UPlanSApo 40×/1.25 NA silicone-immersion objective were used exclusively on the STELLARIS and SPE, respectively. Time-lapse imaging was also performed on a Zeiss LSM900 confocal microscope equipped with a Zeiss Plan-Apochromat 20×/0.8 NA objective. Maximum intensity projections of xy optical sections were generated from confocal z-stacks using Fiji.

### Histology

Third-instar wandering larvae or 1–1.5 h APF pupae were dissected in phosphate-buffered saline (PBS) and fixed in 4% paraformaldehyde (PFA) for 5 min on ice, followed by 20 min at room temperature. Samples were washed three times with PBT (PBS containing 0.1% Triton X-100) for 20 min each, and then blocked with PBTn (5% donkey serum in PBT) for 30 min. For immunostaining, samples were incubated overnight at 4°C with primary antibodies diluted in PBTn. After four 30-min washes with PBT, samples were re-blocked in PBTn for 30 minutes and then incubated with secondary antibodies for 2 h at room temperature. Following four additional 30-min washes with PBT, the samples were mounted in an anti-fade mounting medium composed of 70% glycerol (Sigma, #G5516) and 0.2% n-propyl gallate (KANTO CHEMICAL, #32465-31). For immunostaining for *ex vivo*-cultured wing discs, samples were prepared as described above in the “Sample preparation for *ex vivo* culture of wing discs” section. Cultured wing discs were fixed with 4% PFA for 20 min at room temperature. The subsequent procedures were performed using Terasaki plates (Greiner Bio-One, #654102). Fixed wing discs were washed six times with PBT for 5 min each and then blocked with PBTn for 30 min. Samples were then incubated overnight at 4°C with primary antibodies diluted in PBTn. After eight 5-min washes with PBT, samples were re-blocked in PBTn for 30 min and incubated with secondary antibodies for 2 h at room temperature. Following eight additional 5-min washes with PBT, samples were mounted in an anti-fade mounting medium. For anti-Collagen IV immunostaining, Triton X-100 was not included, and immunostaining was performed in PBS only. All other steps were performed as described above. For detailed observation of the interior of the wing disc, samples were pre-cut using a Feather double-edged razor before imaging. Images were acquired using a Zeiss LSM900 confocal microscope equipped with a Plan-Apochromat 20x/0.8 NA objective or a Zeiss Plan-Apochromat 40x/1.4 NA oil-immersion objective, as well as a Leica STELLARIS confocal microscope equipped with a Leica HC PL APO 20x/0.75 NA objective. Primary antibodies used are as follows; rat anti-GFP antibody (Nacalai Tesque, #04404-26, 1:1000), rabbit anti-GFP antibody (Invitrogen, #A6455, 1:500), rabbit anti-Phospho-Myosin Light Chain 2 (Ser19) antibody (CST, #3671, 1:100), rabbit anti-Zfh1 antibody (R. Lehmann, 1:1000), rabbit anti-DsRed antibody (Clontech, # 632496, 1:500), mouse anti-Gasp antibody (DSHB, #2A12, 1:100), anti-Mmp1 antibody (DSHB, 1:10 from 1:1:1 cocktail of 3A6B4, 3B8D12 and 5H7B11) (PMID: 25224221), and Rabbit anti-Collagen IV antibody (1:100) (PMID: 40570847). Secondary antibodies used are as follows; Goat anti-rabbit Alexa 488 (Invitrogen, #A11034, 1:500), Goat anti-rabbit Alexa 546 (Invitrogen, #A11035, 1:500), Goat anti-rabbit Alexa 647 (Invitrogen, #A21245, 1:500), Goat anti-rat Alexa 488 (Invitrogen, #A11006, 1:500), Goat anti-mouse Alexa 546 (Invitrogen, #A11018, 1:500), Goat anti-mouse Alexa 647 (Invitrogen, #A21237, 1:500). DAPI (Sigma-Aldrich, #D9542, 1:1000) and Phalloidin-647 (Invitrogen, #A22287, 1:400) were applied together with the secondary antibodies.

### Quantification of the final wing disc shape

Images of wing discs cultured *ex vivo* for 24 h were acquired using either a Leica M205C or a Leica M165FC stereomicroscope equipped with a Leica DFC7000T camera, or a Leica SP5, a Leica SPE, or a Leica STELLARIS confocal microscope each equipped with a Leica HC PL APO 10x/0.40 NA objective. The total numbers of wing discs that remained concave or transitioned to convex shape were quantified, and the resulting data were analyzed using RStudio.

### Quantification of phosphorylated MRLC (p-MRLC) signals

To quantify p-MRLC levels, images of wing discs dissected at the third-instar wandering stage or cultured for 12–13 h after dissection, just before the concave-to-convex transition, were acquired using a Zeiss LSM900 confocal microscope equipped with a Zeiss Plan-Apochromat 40x/1.4 NA oil-immersion objective, collecting z-stacks with a total depth of 15 µm. The mean intensity of p-MRLC signals within the hinge region, marked by *30A-Gal4*-driven GFP, was quantified separately for the apical and basal regions using Fiji. The resulting data were analyzed using RStudio.

### Quantification of the Collagen IV-disassembled area on the basal side of wing discs

To compare the extent of Collagen IV disassembly on the basal side, images of wing discs cultured *ex vivo* for 0–1 h or 10–16 h, just before the concave-to-convex transition, were acquired using a Leica STELLARIS confocal microscope equipped with a Leica HC PL APO 20x/0.75 NA objective. The total area of Collagen IV disassembly on the basal side was measured from Z-stack confocal images after maximum-intensity projection and binarization using Fiji. The resulting data were analyzed using RStudio.

### Quantification of the Collagen IV-disassembled area in the notum region of wing discs

To compare the extent of Collagen IV disassembly within the notum, wing discs were cultured *ex vivo* for 10–16 h, just before the concave-to-convex transition. Images were then acquired at the confocal planes where the *viking::GFP* knock-in reporter or anti-Collagen IV antibody staining was most clearly visible, using either a Leica STELLARIS confocal microscope equipped with a Leica HC PL APO 20x/0.75 NA objective or a Zeiss LSM900 confocal microscope equipped with a Zeiss Plan-Apochromat 20x/0.8 NA objective. The total area of Collagen IV disassembly in the notum was measured from Z-stack confocal images after binarization using Fiji, and the resulting data were analyzed using RStudio.

### Quantification of Collagen IV intensity in the pouch region of wing discs

Wing discs were cultured *ex vivo* for 10–16 h, just before the concave-to-convex transition. Images were acquired at the confocal plane corresponding to the largest pouch region within the z-stack using a Leica STELLARIS confocal microscope equipped with a Leica HC PL APO 20x/0.75 NA CS2 objective. The mean intensity of Collagen IV signal within the pouch region was quantified using Fiji, and the resulting data were analyzed using RStudio.

### Quantification of Mmp1-positive puncta

To quantify Mmp1-positive puncta, images of wing discs cultured *ex vivo* for 10–14 h, just before the concave-to-convex transition, were acquired at the confocal plane where Mmp1-positive puncta surrounding Zfh1-positive myoblast nuclei were most abundant, using a Zeiss LSM900 confocal microscope equipped with a Zeiss Plan-Apochromat 40x/1.4 NA oil-immersion objective. The numbers of Mmp1-positive puncta and myoblasts were manually counted using Fiji, and the resulting data were analyzed using RStudio.

### Quantification of Mmp2::GFP intensity in myoblasts

To quantify Mmp2::GFP intensity, images of wing discs cultured *ex vivo* for 10–14 h, just before the concave-to-convex transition, were acquired at the confocal plane where the *Mmp2-GFP* knock-in reporter and Zfh1-positive myoblasts were most clearly visible, using a Zeiss LSM900 confocal microscope equipped with a Zeiss Plan-Apochromat 40x/1.4 NA oil-immersion objective. The mean intensity of Mmp2::GFP signals within Zfh1-positive myoblasts was quantified using Fiji, and the resulting data were analyzed using RStudio.

### Statistical analysis

Statistical analyses were performed, and dot plots and bar graphs were generated using RStudio. Significance levels are indicated as follows: * P<0.05, ** P<0.01, *** P<0.001, and **** P<0.0001.

## Author Contributions

**Conceptualization:** Chigusa Hinata, Shizue Ohsawa.

**Data curation:** Chigusa Hinata.

**Formal analysis:** Chigusa Hinata, Hirotatsu Nakagawa.

**Funding acquisition:** Chigusa Hinata, Shizue Ohsawa.

**Investigation:** Chigusa Hinata, Hirotatsu Nakagawa, Katsuya Nozaki.

**Methodology:** Chigusa Hinata, Shigeaki Nonaka, Yoshikatsu Sato.

**Project administration:** Shizue Ohsawa.

**Resources:** Shigeaki Nonaka, Shizue Ohsawa.

**Supervision:** Shizue Ohsawa.

**Visualization:** Chigusa Hinata, Shizue Ohsawa.

**Writing – original draft:** Chigusa Hinata, Shizue Ohsawa.

**Writing – review & editing:** Chigusa Hinata, Hirotatsu Nakagawa, Katsuya Nozaki, Shigeaki Nonaka, Yoshikatsu Sato, Shizue Ohsawa.

## Acknowledgments

We thank Megumi Nakayama, Emi Maekawa, Keisuke Ikawa, Masayoshi Hayashi, Hiroshi Hosoi, and Takehiro Esaka for discussions, Mina Hoshino, Tomoko Furukawa, and Sayako Suzuki for technical support. We also thank L.S. Shashidhara, K VijayRaghavan, Dirk Bohmann, Barry J. Thompson, Masayuki Miura, J.C. Pastor-Pareja, Ruth Lehmann, the Bloomington *Drosophila* Stock Center (Indiana), the Vienna *Drosophila* Resource Center (Vienna), the National Institute of Genetics Stock Center (Mishima), and the *Drosophila* Genomics and Genetic Resources (Kyoto) for fly stocks. This work was supported in part by JSPS KAKENHI for Transformative Research Area (A) (Grant Nos. JP20H05945 to SO, JP24K21967 to SO), the Scientific Research (B) (Grant No. JP22H02616 to SO), Challenging Exploratory Research (Grant No. JP21K19257 to SO), and Grant-in-Aid for Transformative Research Areas-Platforms for Advanced Technologies, Research Resources “Advanced Bioimaging Support” (Grant No. JP22H04926 to SO), and Grant-in-Aid for JSPS Research Fellows (Grant No. JP24KJ1263 to CH), and Japan Science and Technology Agency (Moonshot Research & Development: Grant Number JPMJPS2022 to SO).

## Supporting information

**S1 Movie. Time-lapse confocal imaging of the concave-to-convex epithelial shape transition in the ecdysone-treated wing disc using confocal microscopy, related to Figure 1**

A wing disc carrying *ubi-RFP* (magenta) was dissected, cultured in the presence of ecdysone and DW as a solvent control, and subjected to time-lapse imaging at 15-min intervals using a Leica STELLARIS confocal microscope. See *Methods* for details.

**S2 Movie. Time-lapse light-sheet imaging of the concave-to-convex epithelial shape transition in the ecdysone-treated wing disc, related to Figure 1**

A wing disc expressing *nls-SCAT3* (green) under the control of *Tub-Gal4* was dissected, cultured in the presence of ecdysone, and subjected to time-lapse imaging at 5-min intervals using a Zeiss Lightsheet Z.1 microscope. The left panel shows the three-dimensional shape of the wing disc, and the right panel shows the corresponding x–z cross-section. See *Methods* for details.

**S3 Movie. Time-lapse confocal imaging of the ecdysone-treated wing disc reveals inhibition of the concave-to-convex epithelial shape transition by the ROCK inhibitor Y-27632, related to Figure 1**

A wing disc carrying *ubi-RFP* (magenta) was dissected, cultured in the presence of ecdysone and the ROCK inhibitor Y-27632, and subjected to time-lapse imaging at 15-min intervals using a Leica STELLARIS confocal microscope. See *Methods* for details.

**S4 Movie. Time-lapse confocal imaging of the concave-to-convex epithelial shape transition in the ecdysone-treated wing disc under DMSO control, related to Figure 2**

A wing disc carrying *ubi-RFP* (magenta) was dissected, cultured in the presence of ecdysone and DMSO, and subjected to time-lapse imaging at 15-min intervals using a Leica STELLARIS confocal microscope. See *Methods* for details.

**S5 Movie. Time-lapse confocal imaging of the ecdysone-treated wing disc reveals inhibition of the concave-to-convex epithelial shape transition by the MMP inhibitor GM6001, related to Figure 2**

A wing disc carrying *ubi-RFP* (magenta) was dissected, cultured in the presence of ecdysone and the MMP inhibitor GM6001, and subjected to time-lapse imaging at 15-min intervals using a Leica STELLARIS confocal microscope. Orange arrowheads indicate the same anatomical position in the disc, which remains concave throughout the imaging period. See *Methods* for details.

**S6 Movie. Time-lapse confocal imaging of Collagen IV in an agarose-embedded, ecdysone-treated wing disc, related to Figure 3**

A wing disc carrying *viking::GFP* (pseudocolor) and *ubi-RFP* (white) was dissected, embedded in agarose gel, cultured in the presence of ecdysone, and subjected to time-lapse imaging at 12-min intervals using a Zeiss LSM900 confocal microscope. See *Methods* for details.

**S7 Movie. Time-lapse confocal imaging of Perlecan in an agarose-embedded, ecdysone-treated wing disc, related to Figure S3**

A wing disc carrying *trol-GFP* (pseudocolor) and *ubi-RFP* (white) was dissected, embedded in agarose gel, cultured in the presence of ecdysone, and subjected to time-lapse imaging at 15-min intervals using a Leica STELLARIS confocal microscope. See *Methods* for details.

**S8 Movie. Time-lapse confocal imaging of the migration of non-epithelial cells in association with Collagen IV disassembly in the ecdysone-treated wing disc, related to Figure S3**

A wing disc carrying *viking::GFP* (green) and *ubi-RFP* (magenta) was dissected, mounted in an agarose gel frame, cultured in the presence of ecdysone, and subjected to time-lapse imaging at 12-min intervals using a Leica SPE confocal microscope. Orange arrowheads indicate the non-epithelial cells that migrate in association with Collagen IV disassembly. See *Methods* for details.

**S9 Movie. Time-lapse confocal imaging of Collagen IV and myoblast dynamics in an agarose-embedded, ecdysone-treated wing disc, related to Figure 3**

A wing disc carrying *viking::GFP* (green) and expressing *RedStinger* (magenta) under the control of *1151-Gal4* was dissected, embedded in agarose gel, cultured in the presence of ecdysone, and subjected to time-lapse imaging at 10-min intervals using a Leica STELLARIS confocal microscope. See *Methods* for details.

**S10 Movie. Time-lapse confocal imaging of Collagen IV and tracheal cell dynamics in an agarose-embedded, ecdysone-treated wing disc, related to Figure 3**

A wing disc carrying *viking::GFP* (green) and expressing *RedStinger* (magenta) under the control of *btl-Gal4* was dissected, embedded in agarose gel, cultured in the presence of ecdysone, and subjected to time-lapse imaging at 10-min intervals using a Leica STELLARIS confocal microscope. See *Methods* for details.

**S11 Movie. Time-lapse light-sheet imaging of the epithelial shape transition and Collagen IV disassembly in the ecdysone-treated wing disc, related to Figure S4** A wing disc carrying *viking::GFP* (green) and *ubi-RFP* (magenta) was dissected, cultured in the presence of ecdysone, and subjected to time-lapse imaging at 20-min intervals using a Zeiss Lightsheet Z.1 microscope. The left panel shows the three-dimensional shape of the wing disc, and the right panel shows the corresponding x–z cross-section. See *Methods* for details.

**S12 Movie. Time-lapse light-sheet imaging reveals that myoblast- and trachea-specific Timp expression inhibits the concave-to-convex epithelial shape transition and Collagen IV disassembly in the ecdysone-treated wing disc, related to Figure S4**

A wing disc carrying *viking::GFP* (green) and *ubi-RFP* (magenta), and expressing *Timp* under the control of *1151-Gal4* and *btl-Gal4* was dissected, cultured in the presence of ecdysone, and subjected to time-lapse imaging at 20-min intervals using a Zeiss Lightsheet Z.1 microscope. The left panel shows the three-dimensional shape of the wing disc, and the right panel shows the corresponding x–z cross-section. See *Methods* for details.

**S13 Movie. Time-lapse confocal imaging reveals that myoblast- and trachea-specific Timp expression inhibits the concave-to-convex epithelial shape transition in the ecdysone-treated wing disc, related to Figure 4**

A wing disc carrying *ubi-RFP* (magenta) and expressing *Timp* under the control of *1151-Gal4* and *btl-Gal4* was dissected, cultured in the presence of ecdysone, and subjected to time-lapse imaging at 12-min intervals using a Leica SPE confocal microscope. See *Methods* for details.

**S14 Movie. Time-lapse confocal imaging reveals that myoblast-specific Timp expression inhibits the concave-to-convex epithelial shape transition in the ecdysone-treated wing disc, related to Figure 4**

A wing disc carrying *ubi-RFP* (magenta) and expressing *Timp* under the control of *1151-Gal4* was dissected, cultured in the presence of ecdysone, and subjected to time-lapse imaging at 12-min intervals using a Leica SPE confocal microscope. See *Methods* for details.

**S15 Movie. Time-lapse confocal imaging reveals that trachea-specific Timp expression inhibits the concave-to-convex epithelial shape transition in the ecdysone-treated wing disc, related to Figure 4**

A wing disc carrying *ubi-RFP* (magenta) and expressing *Timp* under the control of *btl-Gal4* was dissected, cultured in the presence of ecdysone, and subjected to time-lapse imaging at 15-min intervals using a Leica SP5 confocal microscope. See *Methods* for details.

